# A novel vascular endothelial growth factor-trap KP-VR2 with enhanced ligand blocking

**DOI:** 10.1101/2021.02.02.429416

**Authors:** Mi-Ra Kim, Bongkyu kim, Yunsu Na, Jingon Yoo, Se-Ho Kim, In-Ra Seo

## Abstract

Antiangiogenic therapies targeting vascular endothelial growth factor (VEGF)-A have been commonly used in clinics to treat cancers over the past decade. However, their clinical efficacy has been limited, with drawbacks including the acquisition of resistance and activation of compensatory pathways resulting from elevated circulating VEGF-B and placental growth factor (PlGF) levels. Thus, we developed a novel VEGF-Trap, KP-VR2, which can neutralize VEGF-A, VEGF-B, and PlGF to mediate these problems. KP-VR2 consists of two consecutive second Ig-like domains (D2s) of VEGF receptor 1 (VEGFR-1) fused to human IgG1 Fc. KP-VR2 showed more potent decoy activity than the current VEGF-Trap against VEGF and PlGF. Most importantly, two consecutive D2s of VEGFR-1 can generate two putative binding sites, resulting in a significant improvement in binding capacity. These advances resulted in stronger antitumor efficacy in implanted tumor models than aflibercept and bevacizumab. Overall, the results of this study highlight KP-VR2 as a promising therapeutic candidate for further clinical drug development.

## Introduction

Angiogenesis is defined as the neoformation of blood vessels through a multistep mechanism that provides nutrients and oxygen to tissues, allowing the discharge of waste products [1]. This process works in physiological conditions, such as wound healing, embryogenesis, and inflammation, but is also crucial for pathological conditions like cancer [2]. The vascular endothelial growth factor (VEGF) is considered the most important growth factor and is a well-characterized contributor to angiogenesis [3].

VEGF represents the most important and widely studied proangiogenic factor family, and it is composed of five growth factors named VEGF-A, VEGF-B, VEGF-C, VEGF-D, and placental growth factor (PlGF) [4–6]. VEGFs play their role in angiogenesis by binding to three different receptors located on the cell membrane: VEGFR-1 (Flt-1), VEGFR-2 (Flk/KDR), and VEGFR-3 (Flt-4) [7]. VEGF-A is the most important regulator in human physiological and pathological angiogenesis, and it is related to a poor prognosis in several cancers [8]. VEGF-A may interact with both VEGFR-1 and VEGFR-2, but due to the limited kinase activity of VEGFR-1, the active kinase activity of VEGFR-2 makes it the most important effector of VEGF-A downstream signaling [9]. VEGF-B, which has common structural homology with VEGF-A and whose activity is mediated by interaction with VEGFR-1, plays a role in tumorigenesis and blood vessel survival under stress conditions [10]. VEGF-C and VEGF-D, which bind VEGFR-3, are involved in lymphangiogenesis [11].

PlGF is another important growth factor that directly regulates vessel growth and maturation by affecting endothelial cells and indirectly regulates these by recruiting proangiogenic cell types. PlGF shares structural homology with VEGF-A and stimulates angiogenesis through interaction with VEGFR-1 [12–15]. Unlike VEGF-A, which binds to both VEGFR-1 and VEGFR-2, PlGF binds to VEGFR-1 but not to VEGFR-2 [7].

Aflibercept, also known as ziv-aflibercept or VEGF-Trap (Eylea^®^ and Zaltrap^®^, Regeneron Pharmaceuticals, Tarrytown, NY, USA and Sanofi-Aventis, Bridgewater, NJ, USA, respectively), has been approved by the US Food and Drug Administration (FDA) for the treatment of macular degeneration and metastatic colorectal cancer [16]. Aflibercept is a recombinant fusion protein consisting of the second immunoglobulin (Ig) domain of VEGFR-1 and the third Ig domain of VEGFR-2 fused to human IgG1 Fc. It exhibits affinity for VEGF-A, VEGF-B, and PlGF [3, 17].

The VEGF-Trap bevacizumab (Genentech, Inc., San Francisco, CA, USA) is a VEGF blocking monoclonal antibody and has two ligand binding sites. However, bevacizumab binds VEGF-A but not VEGF-B or PlGF.

To address the limitations of the current VEGF-Trap, we designed a new structure of VEGF-Trap KP-VR2 to consist of two consecutive D2s of VEGFR-1 fused to the Fc region of human IgG1. Considering the main role of VEGFR-1 D2 and the minor role of VEGFR-2 D3 in the binding of VEGF [18], we generated KP-VR2 by replacing VEGFR-2 D3 with VEGFR-1 D2 in aflibercept. Thus, two consecutive D2s of VEGFR-1 create two putative ligand binding sites, resulting in improved decoy efficiency compared to aflibercept with one binding site. In addition, KP-VR2 exhibits a high affinity for PlGF, to which bevacizumab cannot bind. This improvement was demonstrated by binding enzyme-linked immunosorbent assay (ELISA), human umbilical vein endothelial cells (HUVECs) migration inhibition, and tumor xenograft models.

## Methods

## Results

### Construction and Expression of KP-VR2

We designed a novel VEGF decoy receptor fusion protein that consists of the two consecutive D2s of VEGFR-1 to create the putative two binding sites for ligands, resulting in greatly improved decoy efficiency (Fig. 1A). The KP-VR2 was produced from CHO cells and purified by protein A affinity chromatography with two additional steps of ion exchange chromatography. The purified KP-VR2 showed ~120 kDa band in a non-reduced SDS-PAGE (Fig. 1B). In western blotting, using an antibody specific to Fc (Fig. 1C), specific bands could be observed for each specified protein at 115 kDa (lane 1, aflibercept Eylea®), 115 kDa (lane 2, aflibercept Zaltrap®), and 120 kDa (lane 3, KP-VR2).

**Figure 1.**
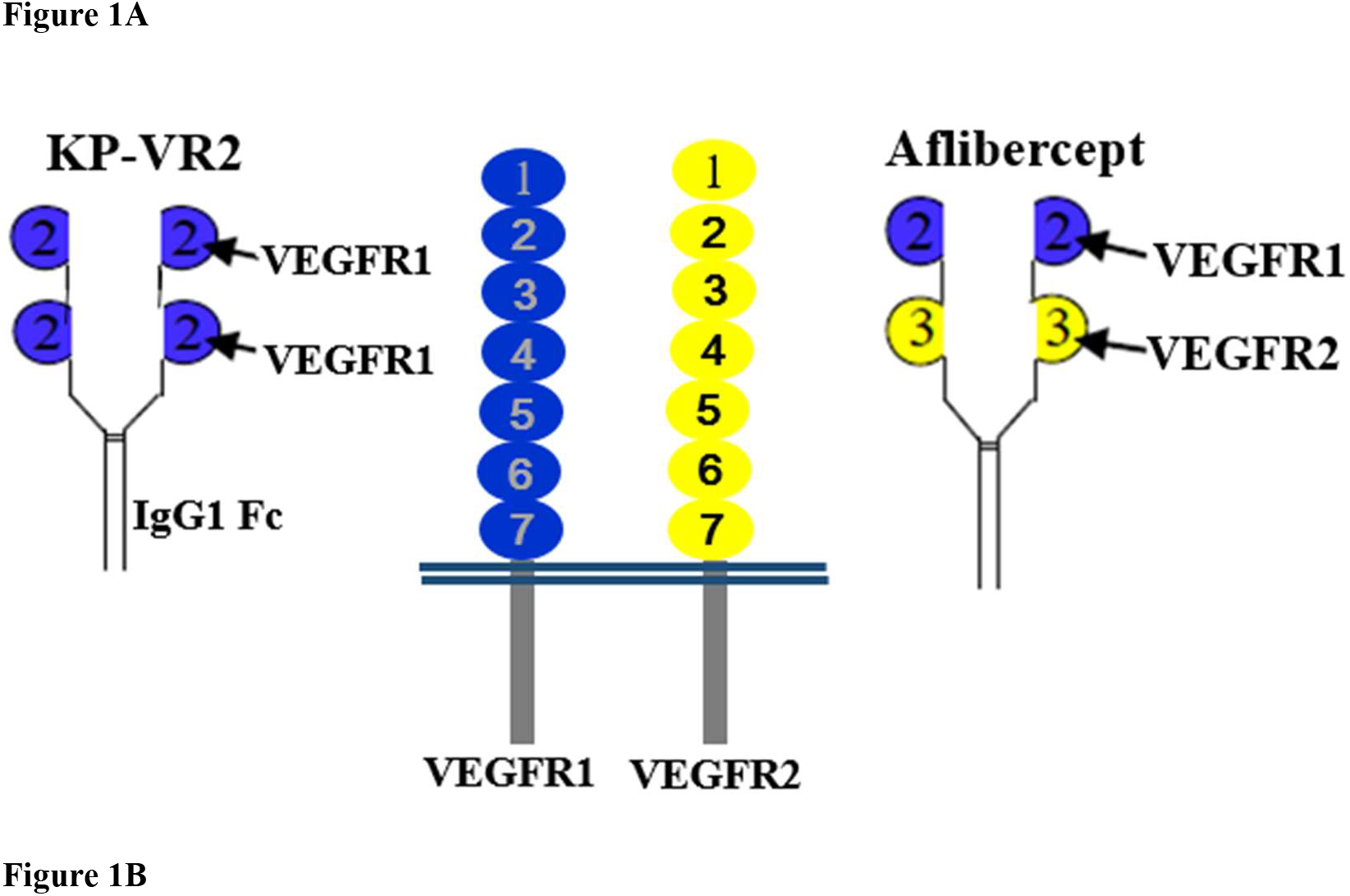
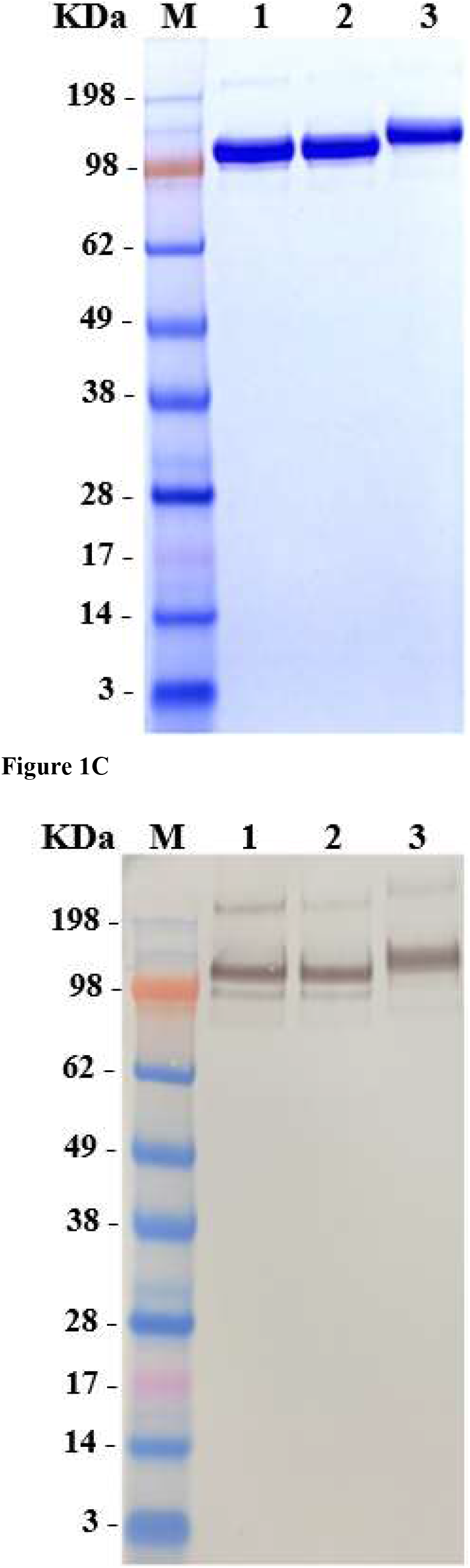
Construction and expression of KP-VR2. Schematic diagram of KP-VR2 and aflibercept. KP-VR2 is composed of VEGFR1 D2–D2-Fc fusion (A). SDS-PAGE analysis of aflibercept and KP-VR2 in non-reduced conditions (B). Western blot analysis of aflibercept and KP-VR2 in non-reduced conditions (C). Lanes 1, 2, and 3 represent aflibercept Eylea®, aflibercept Zaltrap®, and KP-VR2, respectively. Western blot analysis of KP-VR2 and aflibercept was resolved on 4–12% gel under non-reducing conditions, transferred to a polyvinylidene difluoride membrane, and probed with Fc-specific antibody. For comparison, protein MW size marker is shown.

### Comparison of the Binding Ability of KP-VR2, Aflibercept, and Bevacizumab

We compared KP-VR2 to aflibercept and bevacizumab to analyze its ability to bind their ligands *in vitro*. To determine the binding of KP-VR2, aflibercept, and bevacizumab for their ligands, such as VEGF-A_165_, VEGF-A_121_, and PLGF-1, sandwich ELISA assays were performed in which an increasing amount of VEGF-A_165_ and VEGF-A_121_ (from 0.0625 to 256 nM), and PLGF-1 (from 0.122 to 2000 nM) were reacted to immobilized KP-VR2, aflibercept, and bevacizumab. The binding was measured through incubation with Goat anti-Human VEGF antibody and then with peroxidase-conjugated anti-Goat Ig antibody.

The concentration (nM) of 50% of the maximum binding (Bmax_50_) of KP-VR2, aflibercept, and bevacizumab for VEGF-A_165_ was 1.66 nM, 1.69 nM, and 5.16 nM, respectively (Fig. 2A and Table 1). The maximum binding optical density (Bmax_O.D._) of KP-VR2, aflibercept, and bevacizumab for VEGF-A_165_ was O.D. 1.50, 0.89, and 0.91, respectively (Fig. 2A and Table 1).

**Figure 2.**
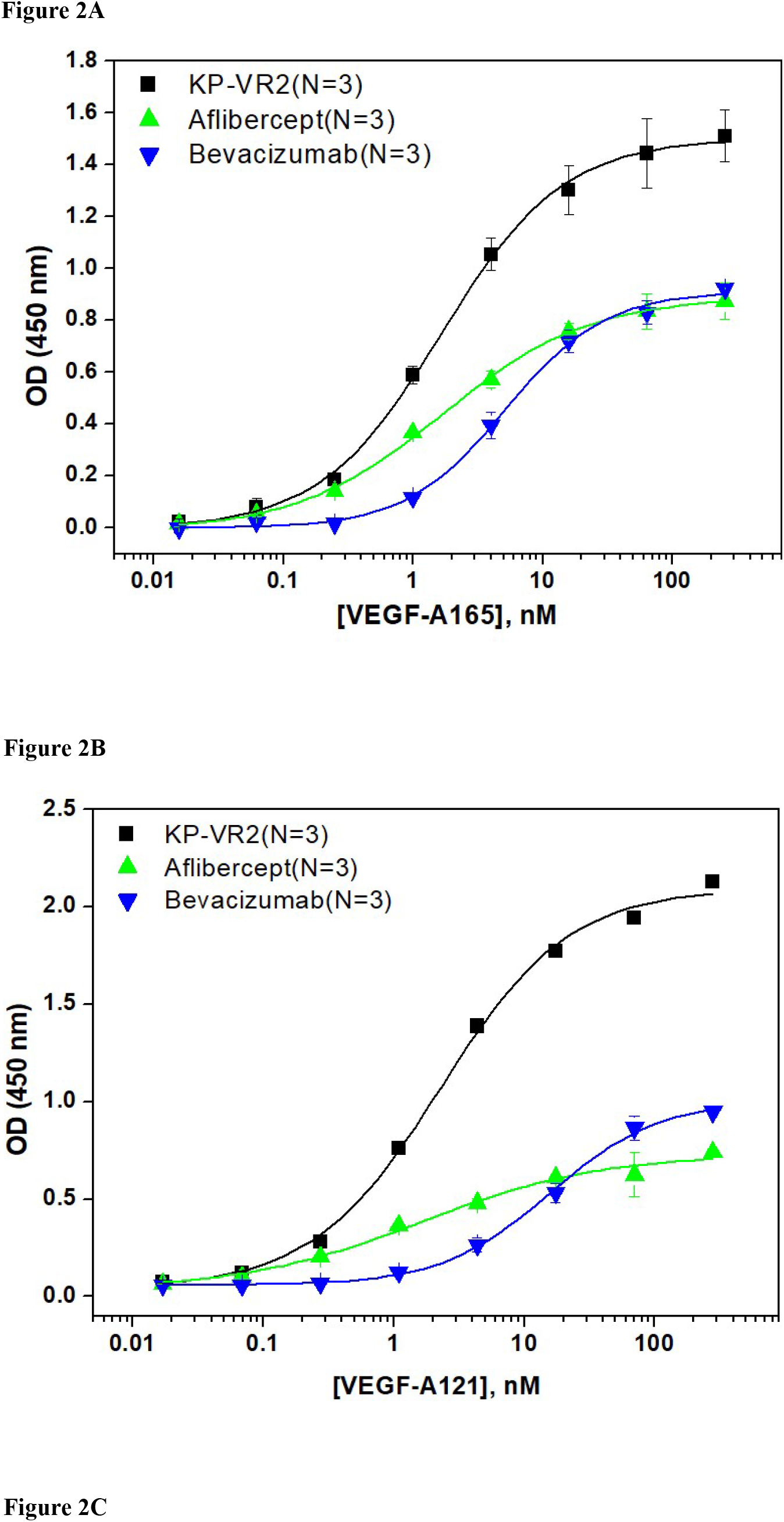
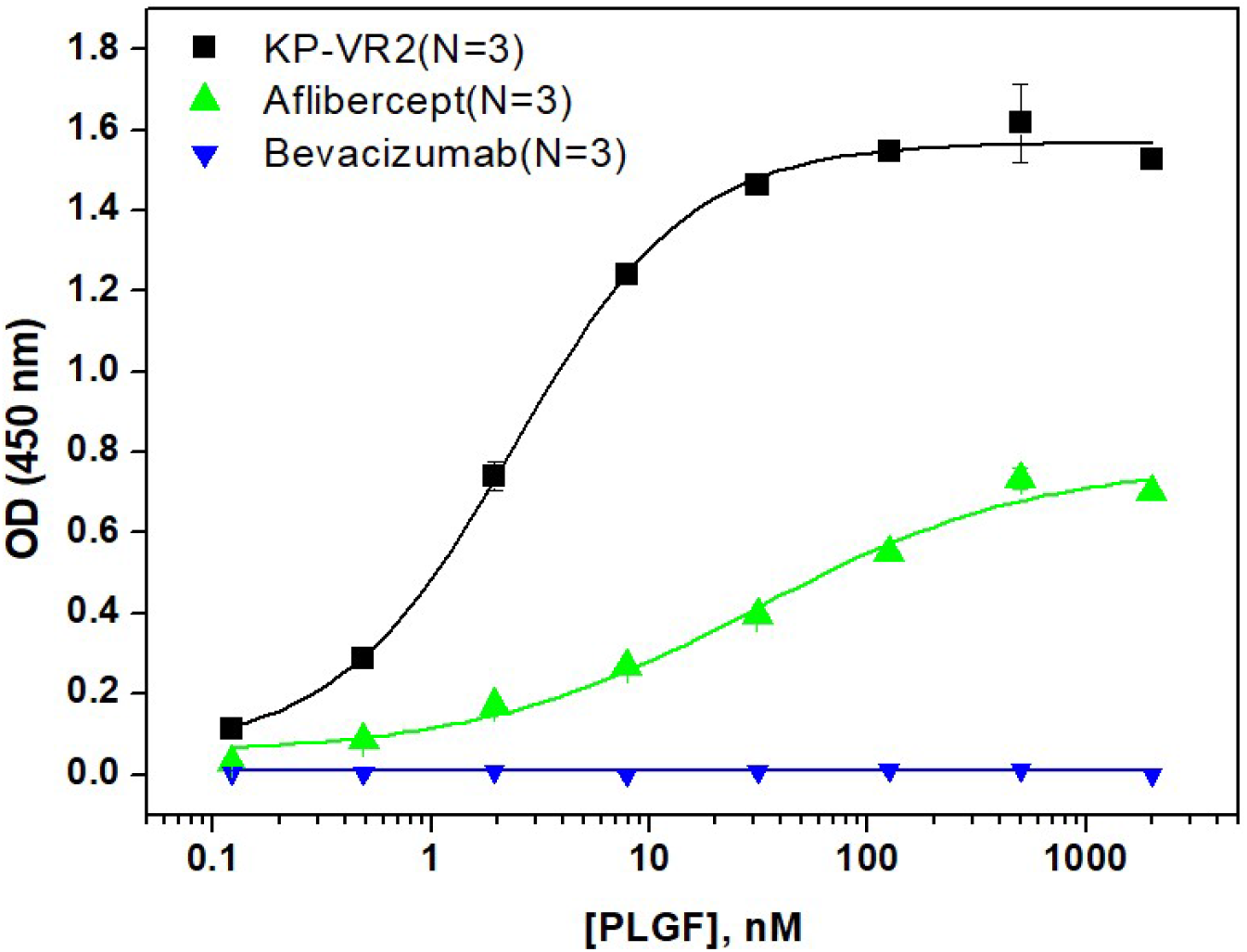
Analysis of binding of KP-VR2, aflibercept, and bevacizumab. Binding abilities of KP-VR2, aflibercept, and bevacizumab to VEGF-A_165_ (A), VEGF-A121 (B), and PlGF-1 (C) are presented. An increasing amount of VEGF-A_165_ and VEGF-A121 1 (from 0.0625 to 256 nM), and PLGF-1 (from 0.1 to 2000 nM) were reacted to immobilized KP-VR2, aflibercept, and bevacizumab. The binding was measured by Goat anti-Human VEGF antibody and then with peroxidase-conjugated anti-Goat Ig antibody. For each group, n = 3. Values are presented as mean ± SD.

**Table 1.** Binding of KP-VR2, aflibercept, and bevacizumab to human VEGF family ligands by ELISA.

| VEGF inhibitor | Ligand | Ligand binding |  |
| --- | --- | --- | --- |
|  |  | Bmax<sub>OD</sub>* | Bmax<sub>50</sub>* (nM) |
| KP-VR2 | VEGF-A<sub>165</sub> | 1.50±0.03 | 1.66±0.15 |
| Aflibercept | VEGF-A<sub>165</sub> | 0.89±0.02 | 1.69±0.15 |
| Bevacizumab | VEGF-A<sub>165</sub> | 0.91±0.02 | 5.16±0.49 |
| KP-VR2 | VEGF-A<sub>121</sub> | 2.10±0.04 | 2.27±0.24 |
| Aflibercept | VEGF-A<sub>121</sub> | 0.73±0.04 | 1.63±0.50 |
| Bevacizumab | VEGF-A<sub>121</sub> | 1.01±0.03 | 16.03±1.59 |
| KP-VR2 | PLGF-1 | 1.57±0.02 | 2.39±0.18 |
| Aflibercept | PLGF-1 | 0.77±0.06 | 30.47±11.46 |
| Bevacizumab | PLGF-1 | - | - |

Bmax<sub>OD</sub> and Bmax<sub>50</sub> represent the maximum binding O.D. and the concentration of ligands at 50% of the maximum binding, respectively

The Bmax_50_ of KP-VR2, aflibercept, and bevacizumab for VEGF-A_121_ was 2.27 nM, 1.63 nM, and 16.03 nM, respectively (Fig. 2B and Table 1). The Bmax_O.D_ of KP-VR2, aflibercept, and bevacizumab for VEGF-A_121_ was O.D 2.10, 0.73, and 1.01, respectively (Fig. 2B and Table 1).

The Bmax_50_ of KP-VR2 and aflibercept for PlGF-1 was 2.39 nM and 30.47 nM, respectively (Fig. 2C and Table 1). The Bmax_O.D_ of KP-VR2 and aflibercept for PlGF-1 was O.D 1.57 and 0.77, respectively (Fig. 2C and Table 1). No binding and bevacizumab binding was detected for PlGF-1 and bevacizumab.

### Comparison of Affinity of KP-VR2 and Aflibercept for VEGF-A_165_ and PLGF-1 by Octet

BLI analysis was performed to assess the binding kinetics of KP-VR2 to VEGF-A and PlGF using a ForteBio Octet System. KP-VR2 and aflibercept were immobilized onto biosensors, which were subsequently dipped into a dilution series of VEGF-A_165_ and PlGF-1. The dynamic reactive process between analytes and ligands was automatically recorded by the Octet control software. The curve and results were analyzed according to the 1:1 binding model (aflibercept) and the 1:2 binding model (KP-VR2). The dynamic association and dissociation curves are shown in Figure 3A to 3B. The kinetics and affinity constants between analytes and ligands are listed in Table 2. Notably, the equilibrium dissociation constant (K_D_) of KP-VR2 for VEGF-A_165_ (<1 pM and 4.75 pM) was similar to that of aflibercept (2.69 pM) (Table 2). KP-VR2 and aflibercept also bound human PlGF-1 with a K_D_ of 0.78 nM and 25.1 nM, respectively. Aflibercept showed a binding affinity to PlGF (1.30 nM).

**Table 2.** Kinetic binding parameters of KP-VR2 and aflibercept to human VEGF family ligands by Octet.

| VEGF inhibitor | Ligand | Kinetic binding parameters |  |  |
| --- | --- | --- | --- | --- |
| | | $K_{on}$ (1/ms) | $K_{dis}$ (1/s) | $K_D$ (M) |
| <sup>1</sup>KP-VR2 | VEGF-A<sub>165</sub> | $3.35 \times 10^4$ | $<1.00 \times 10^{-7}$ | $<1.00 \times 10^{-12}$ |
| <sup>2</sup>KP-VR2 | VEGF-A<sub>165</sub> | $6.44 \times 10^4$ | $3.30 \times 10^{-7}$ | $4.75 \times 10^{-12}$ |
| Aflibercept | VEGF-A<sub>165</sub> | $6.01 \times 10^5$ | $1.64 \times 10^{-6}$ | $2.69 \times 10^{-12}$ |
| <sup>1</sup>KP-VR2 | PLGF-1 | $3.03 \times 10^5$ | $2.36 \times 10^{-4}$ | $7.80 \times 10^{-10}$ |
| <sup>2</sup>KP-VR2 | PLGF-1 | $3.44 \times 10^5$ | $8.65 \times 10^{-3}$ | $2.51 \times 10^{-8}$ |
| Aflibercept | PLGF-1 | $1.74 \times 10^5$ | $2.25 \times 10^{-4}$ | $1.30 \times 10^{-9}$ |

Numbers represent each binding site.
VEGF inhibitor captured on a Protein A-coupled sensor chip

**Figure 3.**
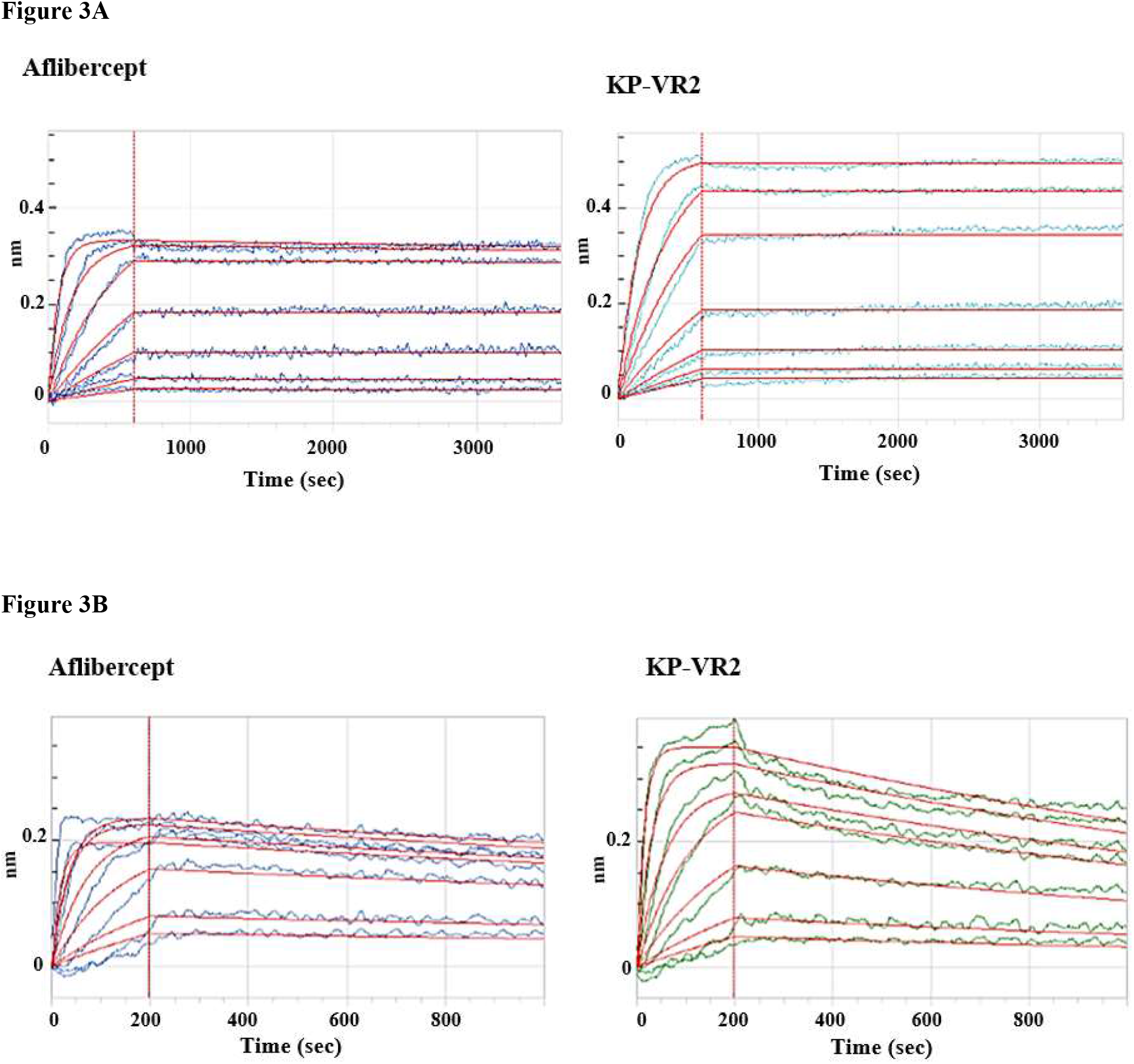
Binding affinities of KP-VR2 and aflibercept for human VEGF-A_165_ and PlGF-1. The interaction between KP-VR and aflibercept with human VEGF-A_165_ and PlGF-1 was measured using Octet. KP-VR2 and aflibercept bound human VEGF-A_165_ with high affinity (*K*_*D*_ = <1 pM and 4.75 pM vs 2.69 pM) (A). KP-VR2 and aflibercept also bound human PlGF-1 with high affinity (*K*_*D*_ = 0.78 nM and 25.1 nM) (B).

### Comparison of KP-VR2, Aflibercept, and Bevacizumab in migration inhibition

We determined whether KP-VR2 binding to VEGF could effectively block the ability of VEGF-induced endothelial cell migration. The inhibitory effect of KP-VR2 was evaluated by endothelial cell invasion and migration assay. KP-VR2, aflibercept, and bevacizumab could completely block VEGF-A_165_-induced HUVEC migration at the concentration of 100 ng/mL. KP-VR2 had a lower half maximal inhibitory concentration (IC_50_) value than aflibercept and bevacizumab (Figure 4B and Table 3) owing to a higher inhibition potency. VEGF induced cell invasion in HUVEC, whereas KP-VR2, aflibercept, and bevacizumab inhibited VEGF-induced cell invasion (Fig. 4A). The blue-colored spot indicates VEGF-induced cell migration (Fig. 4A). KP-VR2 showed higher inhibitory activity than aflibercept and bevacizumab at the same concentration (Fig. 4B). The IC_50_ values of KP-VR2, aflibercept, and bevacizumab were 11 nM, 20 nM, and 21 nM, respectively (Table 3), which reveals that the KP-VR2 has outstanding inhibitory activity against cell invasion.

**Table 3.** IC_50_ of KP-VR2, aflibercept, and bevacizumab in HUVEC migration.

| VEGF inhibitor | Ligand | HUVEC invasion |
| --- | --- | --- |
|  |  | IC<sub>50</sub> (nM) |
| KP-VR2 | VEGF-A<sub>165</sub> | 11.55±0.96 |
| Aflibercept | VEGF-A<sub>165</sub> | 19.93±2.62 |
| Bevacizumab | VEGF-A<sub>165</sub> | 21.15±2.69 |

IC<sub>50</sub> represents half maximal inhibitory concentration.

**Figure 4.**
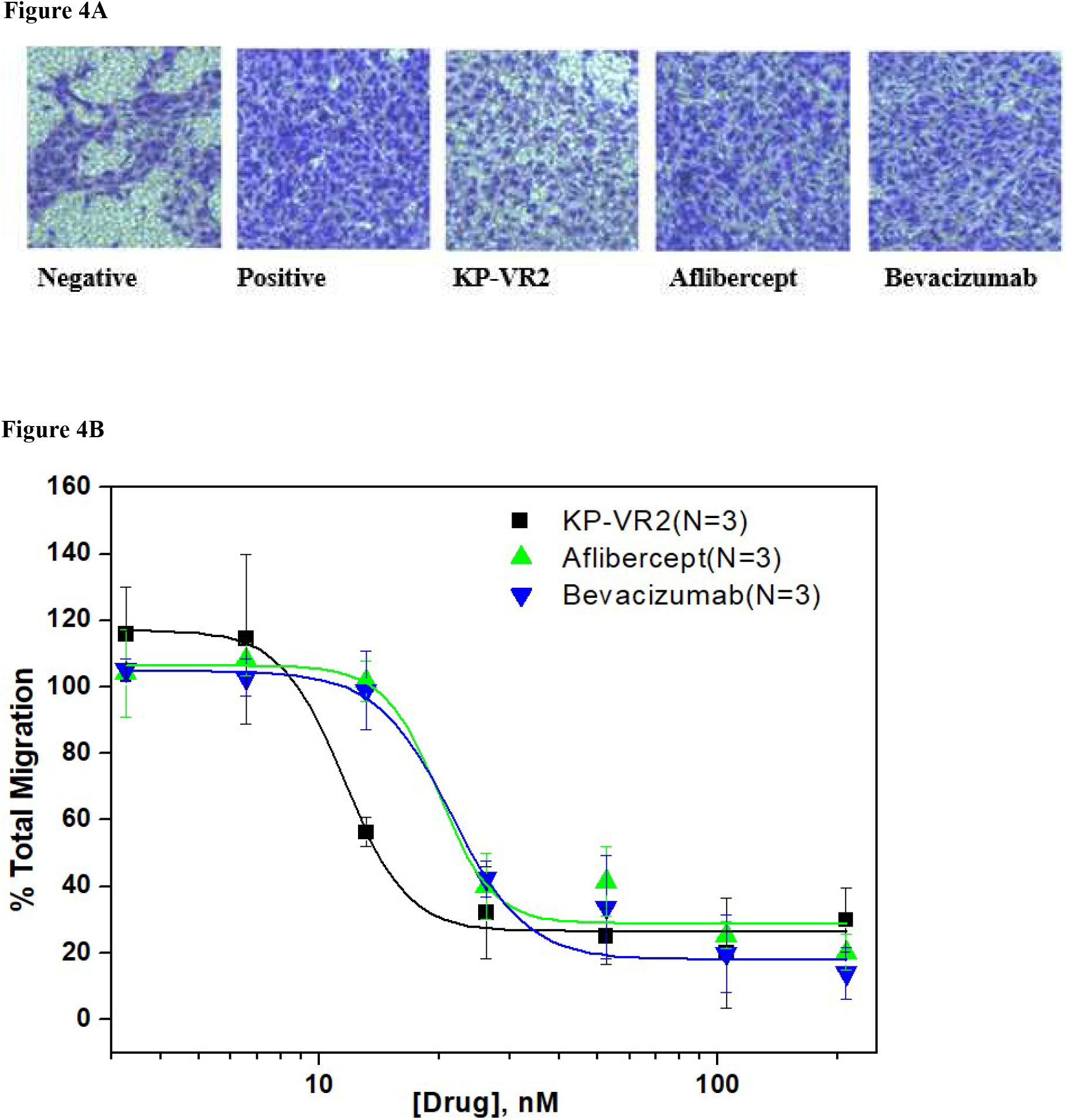
Inhibition of HUVEC migration by KP-VR2, aflibercept, and bevacizumab. Cell invasion assay with HUVEC in the presence of VEGF-A_165_ and VEGF-Traps is presented by image (A) and quantification (B). HUVECs were placed in the upper compartment of the Boyden chamber and allowed to migrate toward basal media containing 0.1% fetal bovine serum with or without VEGF-A_165_ and VEGF-A_165_ mixed with four different concentrations of KP-VR2, aflibercept, and bevacizumab ranging from 3.27 to 209.44 nM. The percentage of total migration (y-axis) was calculated as described in the Experimental Procedures section.

### Comparison of tumor growth inhibition by KP-VR2 and Aflibercept

The inhibition of tumor growth by KP-VR2 and aflibercept was evaluated in a mouse xenograft model. Tumor cells were derived from diverse tissue origins (human LOVO colorectal adenocarcinoma cells, human SKUT-1B mesodermal tumor, and human HT-29 colorectal adenocarcinoma cells).

KP-VR2 significantly inhibited the growth of all three types of tumors (Fig. 5). To evaluate the antitumor effects of KP-VR2, we used the LOVO tumor model and treated them with either KP-VR2 or aflibercept at 1 mg/kg. KP-VR2 treatment resulted in 84% and 88% reduction in tumor volume and weight, whereas aflibercept treatment caused a reduction of 66% and 73%, respectively (Fig. 5A and 5B). We also assessed the antitumor effect of KP-VR2 on established SKUT-1B tumors. These results showed 80% and 51% reduction in tumor volume after 2 mg/kg KP-VR2 and aflibercept treatment, respectively (Fig. 5C and 5D), implying that KP-VR2 is a promising agent against tumors. In the study using HT-29 colorectal adenocarcinoma cells, a 3-fold lower dose KP-VR2 (1 mg/kg) was tested and found to be equally effective at inhibiting tumor growth and weight (Fig. 5E and 5F). Taken together, these results suggest that the improved avidity of KP-VR2 to VEGF-A and PlGF contributes to its higher efficacy in cancer treatment compared to aflibercept.

**Figure 5.**
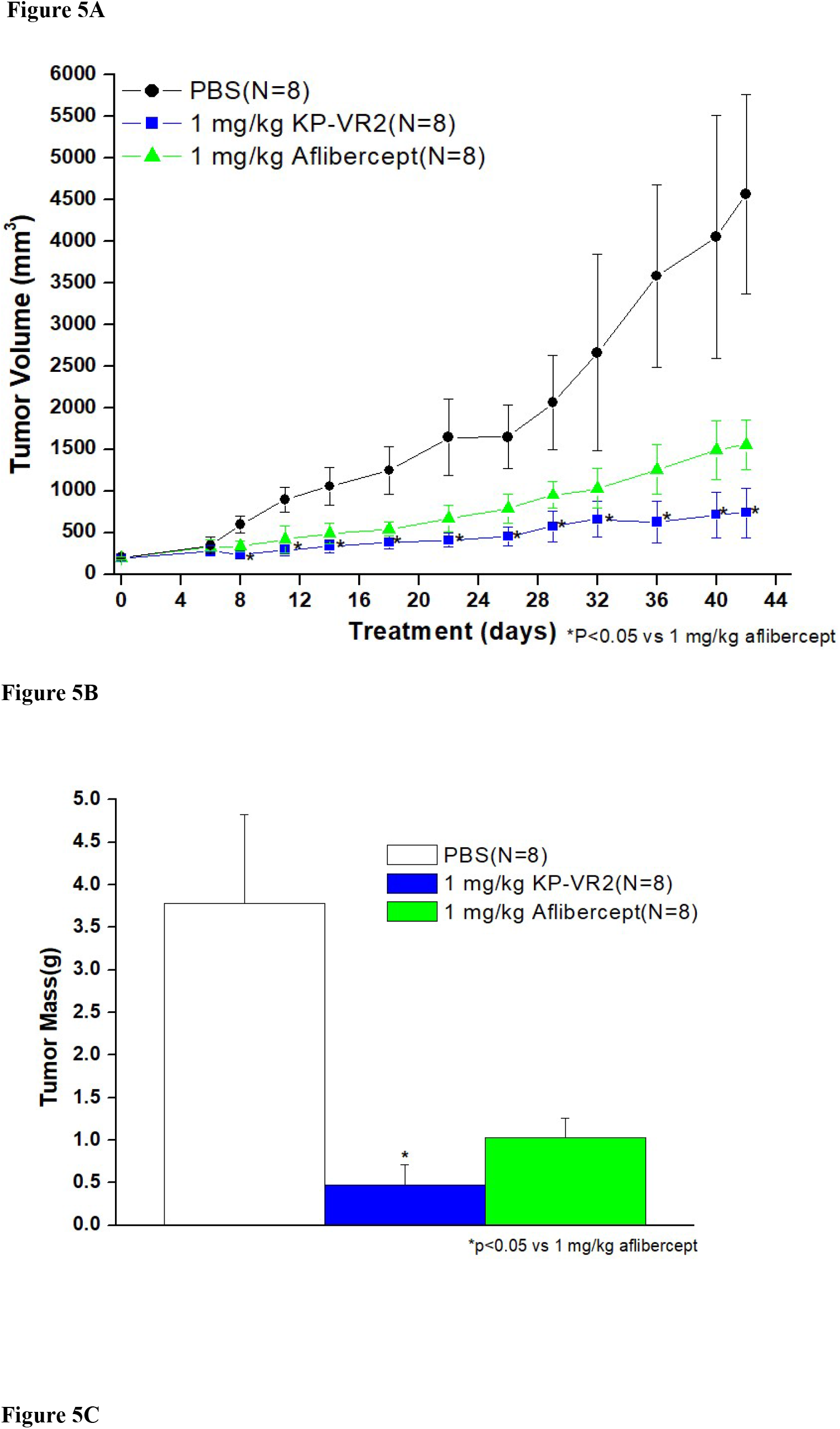
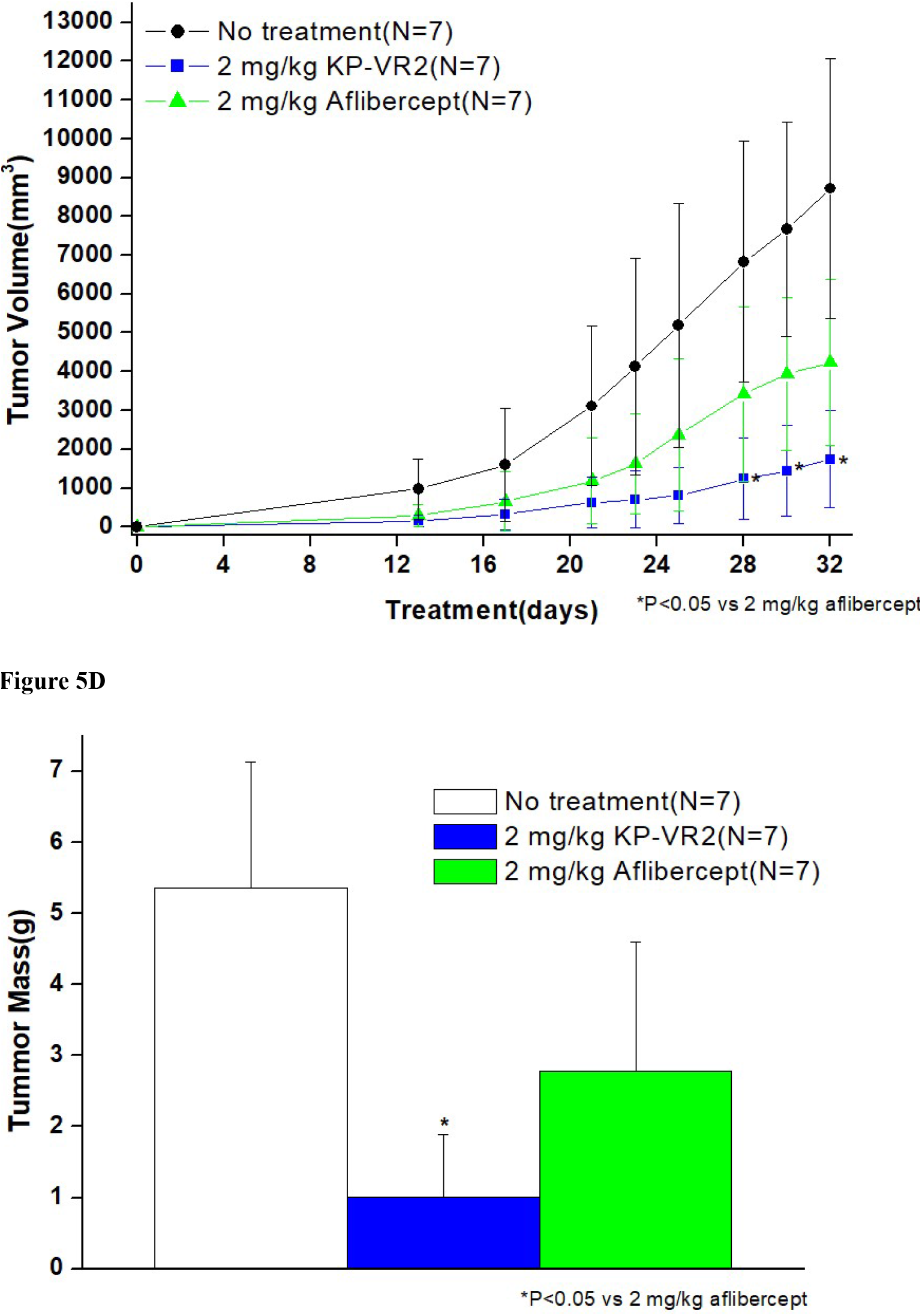
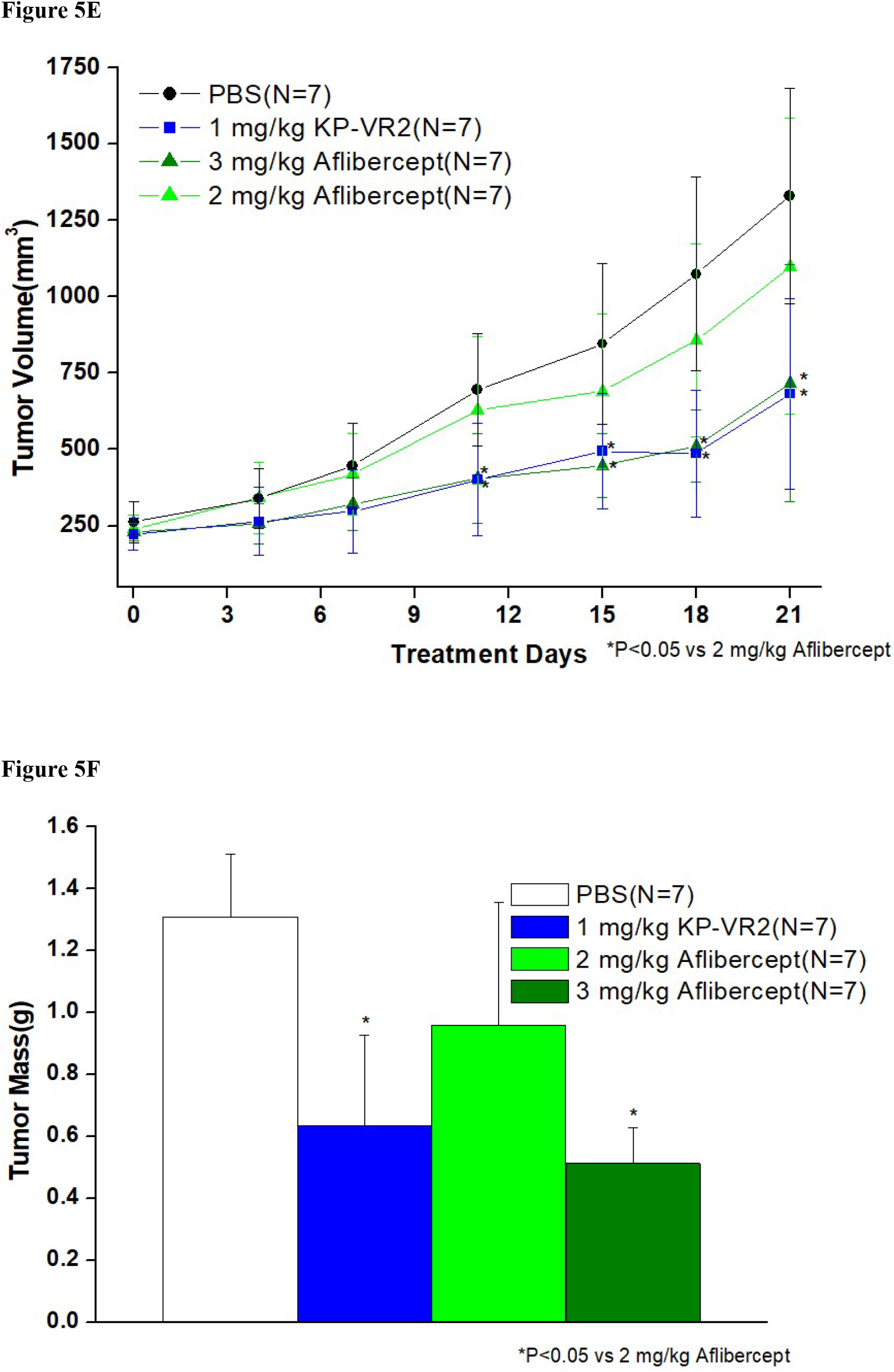
Inhibition of tumor growth in LOVO, SKUT-1B, and HT-29 tumors by KP-VR2 and aflibercept. Tumor growth inhibition induced by KP-VR2 in LOVO, SKUT-1B, and HT-29 models was compared with that of aflibercept. Tumor growth curves were plotted for KP-VR2, aflibercept, and control groups. Tumor volume (A) and mass (B) in LOVO tumor model. Tumor volume (C) and mass (D) in SKUT-1B tumor model. Tumor volume (E) and mass (F) in HT-29 tumor model. * indicates P < 0.05, and all values are mean ± SD.

## Discussion

Angiogenesis continues to be a viable therapeutic target for pathological conditions, including cancer and age-related macular degeneration. The development and investigation of bevacizumab and aflibercept as therapeutic agents has provided a basis for understanding the clinical potential of biological therapies that target angiogenesis.

The validation of VEGF as an important new target against cancer comes from pioneering clinical studies using a humanized monoclonal antibody that binds and blocks VEGF. Because anti-VEGF approaches act by blocking tumor-associated angiogenesis, which appears to be widely required by many different types of tumors, these approaches may prove to be generally useful against a wide assortment of cancers [22]. In addition, pathological angiogenesis seems to contribute to a number of non-neoplastic diseases, such as diabetic retinopathy [23] and psoriasis [24], extending the potential utility of anti-VEGF therapeutics. This highlights the need to optimize anti-VEGF approaches.

KP-VR2 (Fig. 1A) is an Fc fusion protein of two consecutive D2s from VEGFR-1, which was designed on the basis of a previously published report [25], in which the critical role of VEGFR-1 domain 2 was demonstrated to have high affinity binding.

VEGFR-1 has seven Ig-like domains in the extracellular domain (Fig. 1A). Among them, VEGFR-1 D2 is the primary contributor to VEGF-A and PlGF binding. Additionally, residues in VEGFR-1 D3 also participate in the high-affinity binding of VEGF-A and PlGF [26, 27]. Thus, the minimal required domain to bind both VEGF-A and PlGF with high affinity is VEGFR-1 D2–D3.

Although VEGFR-1 binds to VEGF-A and PlGF with higher affinity than VEGFR-2, the development of therapeutic decoy proteins with the VEGFR-1 backbone has proven to be difficult thus far. The major reason behind this is the high pI value of VEGFR-1 owing to the positively charged residues in the VEGFR-1 D3 region. This causes nonspecific extracellular matrix binding and poor pharmacokinetic profiles, leading to a shortened half-life, a subsequent decrease in efficacy, and even toxic side effects [25]. We assume that aflibercept employed VEGFR-2 D2 instead of VEGFR-1 D2 due to the intrinsic problem of VEGFR-1 D2.

However, previous structural analyses have indicated that VEGFR-1 might make greater use of its D2 in contacting VEGF [28]. Therefore, to rule out the problem of VEGFR-1 D3, we generated KP-VR2 consisting of two consecutive D2s of VEGFR-1, demonstrating improved decoy efficiency and dramatic increased avidity for both VEGF and PlGF.

Creating a new decoy receptor fusion protein with an additional binding site has great advantages. Most importantly, the two consecutive D2s of VEGFR-1 mainly consisted of two putative binding sites for ligands, resulting in greatly improved decoy efficiency, as shown in Fig. 2. The combination of high-affinity and improved avidity contributes to making KP-VR2 one of the most potent and efficacious VEGF blockers available. KP-VR2 showed more potent decoy activity against VEGF-A and PlGF than aflibercept. In addition, KP-VR2 showed more potent decoy activity against VEGF-A than bevacizumab, but bevacizumab was unable to bind PlGF. This was evidenced by our *in vitro* experiments demonstrating the strong suppression of proliferation and migration after KP-VR2 treatment. Consistent results were observed *in vivo*, where KP-VR2 showed much stronger antitumor effects in implanted tumor models when judged against aflibercept and bevacizumab.

When equimolar doses of KP-VR2 and aflibercept were compared in the LOVO, SKUT-1B, and HT-29 xenograft models, it was apparent that much higher doses of aflibercept were required to inhibit tumor growth (Fig. 5). When used at the same dose, KP-VR2 showed better efficacy than aflibercept. Furthermore, KP-VR2 and aflibercept have similar circulation times in mice and rats (data not shown). Considering that the HT-29 tumor models are known to be relatively resistant to anti-VEGF therapy [27], these findings suggest the possibility of overcoming resistance to anti-VEGF therapy by concurrent blockade of PlGF with KP-VR2.

An additional advantage is that KP-VR2 is composed entirely of human sequences, minimizing the possibility of immunogenicity in human patients. Additionally, KP-R2 bound all species of VEGF tested, such as human, rabbit, rat, and mouse VEGF (data not shown), rendering it a versatile reagent that can be used in almost any experimental animal model.

Currently, anti-VEGF agents are approved for clinical use in combination with chemotherapy for the treatment of various tumors [22]. FDA approved aflibercept, in combination with leucovorin, 5-fluorouracil, and irinotecan (FOLFIRI), for treating patients after progression with oxaliplatin-containing regimens [2, 16]. This soluble decoy receptor shows one-to-one high-affinity binding to all isoforms of VEGF and PLGF [18, 29]. Aflibercept is also FDA approved for the treatment of age-related macular degeneration (AMD), exhibiting higher efficiency than a single approach for the inhibition of VEGF-A [24].

In this study, KP-VR2 was designed to bind and sequester VEGF as well as PlGF with high affinity and avidity. Like aflibercept, KP-VR2 can be applied to treat various tumors and angiogenic ocular diseases, including AMD, and will be evaluated in future clinical trials. Our results suggest that KP-VR2 is a potent and effective decoy for both VEGF and PlGF. Through the enhanced ligand binding avidity, KP-VR2 inhibited tumor growth and cell migration more effectively than did current VEGF-Traps. The clinical applicability of this novel fusion protein should be further explored through preclinical and clinical studies.

## Experimental Procedures

### Reagents and Materials

Dulbecco’s modified Eagle medium, fetal bovine serum (FBS), RPMI-1640, and trypsin-EDTA were purchased from Gibco (Gaithersburg, MD, USA). F12K and McCoy’s 5A were purchased from ATCC (Rockville, MD, USA). Monoclonal antibodies against VEGFR-2, Erk1/2 phospho-VEGFR-2 (Tyr1175), and phospho-p44/42 MAPK (Erk1/2) (Thr202/Tyr204) were purchased from Cell Signaling Technology (Beverly, MA, USA). Anti-human IgG-peroxidase antibodies were purchased from KPL (Gaithersburg, MD, USA). Human VEGF-A_165_, human VEGF-A_121_, and human PlGF-1 were purchased from R&D Systems (Minneapolis, MN, USA). Aflibercept (Regeneron Pharmaceuticals, Inc., Tarrytown, NY, USA) and bevacizumab (Genentech, Inc., San Francisco, CA, USA) were purchased.

### Cell Lines

We used the following cells in the study: HUVECs (C-2519. Lonza, Walkersville, MD, USA), Chinese hamster ovary suspended cells (CHO-S; A1369601, Gibco, Gaithersburg, MD, USA), LOVO (CCL-229, ATCC), SKUT-1B (HIB-115, ATCC), and HT-29 (HIB-38, ATCC).

### Cell Culture

HUVECs were cultured in EGM-2 medium supplemented with 2% FBS, hydrocortisone, hFGFB, VEGF, R3-IGF, ascorbic acid, hEGF, GA-1000, and heparin. These cells were used to evaluate the effects of KP-VR2 on VEGFR-2.

LOVO, SKUT-1B, and HT-29 cells were grown in F12K, Eagle’s Minimum Essential Medium, and McCoy’s 5A or F12K with 10% FBS, respectively. These cells were allowed to grow to a confluence of approximately 80% and were subcultured 2 or 3 times a week.

### Construction and Expression of KP-VR2

KP-VR2 was created by fusing two consecutive second Ig domains of VEGFR-1 to Fc of human IgG1 (D2-D2-Fc). The KP-VR2 expression construct was transfected into CHO-S cells using Lipofectamine 2000 (Life Technologies) and cultured as previously reported [19]. KP-VR2 was secreted into the culture media and purified by protein A-Sepharose affinity chromatography, followed by two different cation and anion exchange chromatography runs.

The purified KP-VR2 was analyzed by non-reducing sodium dodecyl sulfate-polyacrylamide gel electrophoresis (SDS-PAGE) and western blotting. SDS-PAGE analysis of aflibercept and KP-VR2 was resolved on 4–12% gel under non-reducing conditions.

In the western blot analysis, the KP-VR2 and aflibercept were resolved on 4–12% gel under non-reducing conditions, transferred to a polyvinylidene difluoride membrane, and the bands were probed with human Fc-specific antibody (KPL, Gaithersburg, MD, USA) and horseradish peroxidase-conjugated Goat anti-Human IgG (Seracare, Milford, MA, USA) and a DAB substrate kit (VECTOR Lab, Burlingame, CA, USA).

### Analysis of Binding of KP-VR2, Aflibercept, and Bevacizumab

The binding abilities of KP-VR2, aflibercept, and bevacizumab were measured by ELISA. For this, a 96-well plate was coated with 1–3 nM of KP-VR2, aflibercept, and bevacizumab and blocked with 0.3% bovine serum albumin (BSA) in PBST (Phosphate-Buffered Saline Tween-20). Then, VEGF-A_165_ and VEGF-A_121_ were added in gradually increasing amounts from 0.0625 nM to 256 nM, and PLGF-1 from to 2000 nM. Next, the plate was washed and reacted with the Goat anti-Human VEGF antibody. Then, the plate was washed again and reacted with peroxidase-conjugated anti-Goat Ig antibody. Next, 3,3’,5,5’-tetramethylbenzidine (TMB) solution (Bethyl Laboratories, Montgomery, TX, USA) was added, and, thereafter, absorbance was measured at 450 nm. The data were analyzed by a four-parameter logistic equation (Origin v7).

### Biolayer Interferometry (BLI) Analysis

Binding kinetics were measured on the Octet QK384 System (ForteBio, Pall Life Science, Fremont, CA, USA), which is based on BLI technology.

KP-VR2 and aflibercept were immobilized onto anti-Human Fc biosensors (Pall ForteBio) at a concentration of 20 nM for 600 s, followed by biosensor rinsing in a kinetics buffer that served as a background buffer (KB buffer, Pall ForteBio). After they were balanced to the baseline, the biosensors were dipped into serial dilutions of VEGF and PlGF. The association was measured by the incubation of biosensors in various concentrations (0.3125–20 nM) of VEGF and PlGF for 600 s. Subsequently, dissociation was performed for 3,000 s. The analyses of binding kinetics were evaluated by ForteBio data analysis software using curve fit models of 1:1 binding and 2:1 heterogeneous binding.

### Cell Invasion Assay

The HUVEC invasion assay was performed *in vitro* using a transwell chamber system with 8.0-μm pore polycarbonate filter inserts (Corning Costar Corporation, Cambridge, MA, USA). Briefly, the 8-μm pore inserts were coated with 5 μg/mL collagen (Collagen Type I, Rat Tail, Upstate). HUVECs (7.5 × 10^4^ cells/300 μL) were seeded to the upper wells in EBM-2 medium with 0.1% FBS. EBM-2 medium (750 μL), with 0.1% FBS in the presence of VEGF-A_165_ (350 ng/mL) and KP-VR2, aflibercept, or bevacizumab (0–200 nM), was placed in the lower wells. The cells were incubated at 37 °C for 48 h. After 48 h, the upper collagen-coated surface was wiped off using a cotton swab. Cells that migrated through the filters were fixed, stained with crystal violet, photographed, and counted. The migrated cells were determined using a multi-detection microplate reader at an excitation of 458 nm and emission of 528 nm (BioTek Instrument, Inc., Winooski, VT, USA). The percentage of total migration (y-axis) was calculated as (F_Drug_ - F_Basal_)/ (F_Total_ - F_Basal_) where F_Total_ is fluorescence in the presence of VEGF- A_165_, F_Basal_ is fluorescence in the absence of VEGF- A_165_, and F_Drug_ is fluorescence in the presence of VEGF-A_165_ mixed with drug at a specific molar ratio (x-axis) [20]. The values were analyzed by a four-parameter logistic equation (Origin v7).

### Tumor Growth Inhibition Experiments

Tumor cells (5.0 × 10^6^ cells/mouse) were suspended in PBS and implanted subcutaneously on the shaved right flank of 8–10-week-old male BALB/c mice (Orient Bio Inc.). Tumor-bearing mice were randomized into control and treatment groups (n=7–8 per group).

Then, the mice received an intraperitoneal injection of PBS, KP-VR2, or aflibercept twice weekly during the experiment. The size of the tumor was measured with calipers (tumor volume = length × width × height).

For LOVO and HT-29 models, eight mice bearing tumors of 150–300 mm^3^ were injected intraperitoneally twice a week with 1 mg/kg KP-VR2, 1 and 3 mg/kg aflibercept, and vehicle, 100 μL of PBS. For the SKUT-1B model, the mice were allowed a brief recovery period (one day) and were then injected intraperitoneally twice a week with 2 mg/kg KP-VR2 and aflibercept until the end of the experiment, after which the animals were euthanized and tumors were excised and measured.

### Statistical Analyses

Statistical differences between means were determined by an independent samples t test. The statistical significance was set at *P* < 0.05.

## Data Availability Statement

The data used to support the findings of this study are included within the article.

